# Temporal fluctuations in the brain’s modular architecture during movie-watching

**DOI:** 10.1101/750919

**Authors:** Richard F. Betzel, Lisa Byrge, Farnaz Zamani Esfahlani, Daniel P. Kennedy

## Abstract

Brain networks are flexible and reconfigure over time to support ongoing cognitive processes. However, tracking statistically meaningful reconfigurations across time has proven difficult. This has to do largely with issues related to sampling variability, making instantaneous estimation of network organization difficult, along with increased reliance on task-free (cognitively unconstrained) experimental paradigms, limiting the ability to interpret the origin of changes in network structure over time. Here, we address these challenges using time-varying network analysis in conjunction with a naturalistic viewing paradigm. Specifically, we developed a measure of inter-subject network similarity and used this measure as a coincidence filter to identify synchronous fluctuations in network organization across individuals. Applied to movie-watching data, we found that periods of high inter-subject similarity coincided with reductions in network modularity and increased connectivity between cognitive systems. In contrast, low inter-subject similarity was associated with increased system segregation and more rest-like architectures. We then used a data-driven approach to uncover clusters of functional connections that follow similar trajectories over time and are more strongly correlated during movie-watching than at rest. Finally, we show that synchronous fluctuations in network architecture over time can be linked to a subset of features in the movie. Our findings link dynamic fluctuations in network integration and segregation to patterns of intersubject similarity, and suggest that moment-to-moment fluctuations in FC reflect shared cognitive processing across individuals.

## INTRODUCTION

Cognitive processes are supported by the transient coupling and uncoupling of activity between distant brain regions [1, 2]. These patterns can be modeled as networks, whose nodes and edges represent regions and their pair-wise functional connectivity (FC) [3]. Brain networks can then be investigated using methodology from network science [4, 5], a discipline that provides both a conceptual and mathematical framework for investigating the architecture and dynamics of real-world networks.

Most FC analyses have focused on intrinsic connectivity, which can be reconstructed from brain activity recorded during rest, i.e. task-free conditions [6]. One of the most salient features of resting FC is its modular organization, which refers to the decomposability of the brain into segregated sub-networks called “modules” or “communities” [7, 8]. The boundaries of these modules closely recapitulate patterns of task-evoked activity [9], are highly replicable [10–12], and are thought to engender specialized brain function [13].

Interestingly, the level of segregation in the brain can be modulated. One contributing factor is cognitive state; performing cognitively-demanding tasks requires coordination between modules that at rest are functionally isolated from one another [14]. This constraint results in increased inter-modular FC, which effectively decreases segregation. This phenomenon is general, and has been reported across a wide range of tasks [15–18].

Another factor that can modulate segregation is time. So-called “time-varying” FC can be estimated by partitioning a scan session into a series of windows and estimating FC separately for each window, making it possible to track changes in network architecture over short timescales [19, 20]. Using this approach, many studies have reported temporal fluctuations in the brain’s level of segregation, suggesting that modules transiently couple and uncouple with one another over time [21–23]. This has given rise to hypothesis that the brain alternates between segregated and integrated states, reflecting periods of local, specialized information processing and inter-modular information transfer, respectively [17, 23– 25]

Unlike task FC, where variation in segregation can be attributed to task-related cognitive processes, time-varying FC analysis is usually applied to resting-state datasets in which subjects’ cognitive state is not experimentally controlled. This makes it difficult to ascertain whether fluctuations in segregation are driven by coincident fluctuations in neurophysiological variables like drowsiness or attention [26, 27], internal cognitive processes [28–30], or merely reflect sampling variability around a temporally stationary pattern of FC [31, 32].

Indeed, disambiguating meaningful fluctuations in time-varying FC from statistical noise during rest [33] and tasks [34] remains an open challenge. Here, we propose addressing this question using naturalistic imaging, which involves presenting subjects with short videos (though for consistency with the literature we refer to these as “movies” throughout this manuscript) during the course of a scan session. Movie-watching can be considered a state situated somewhere between that of rest and task, in which subjects are presented with identical stimuli (akin to task) but are not instructed to respond in any particular way (akin to rest).

Recently, movie-watching paradigms have become popular within the neuroimaging [35–38] and network neuroscience communities [39–42], where FC estimated from movie-watching exhibits greater test-retest reliability [43], benefits from reduced in-scanner head motion [44, 45], and enhances the identifiability of individual subjects [46] compared to rest. Additionally, movies feature correlated categories that better match the statistical properties of our day-to-day experiences. For example, rather than viewing a disembodied human face, faces that appear in a movie are usually accompanied by speech, movement, narrative, and context. Despite this, the underlying principles that govern fluctuations in FC over short timescales during movie-watching remain poorly understood.

To address this question, we estimate time-varying FC by applying sliding-window analysis to movie-watching data. We take advantage of the fact that subjects observe the same time-locked stimuli to focus on moments in time when FC across subjects coalesces into similar patterns. We develop a statistical test for identifying those periods, and discover that high inter-subject similarity is linked to decreases in segregation, as indexed by the modularity measure. Next, we show that time-varying FC during movie-watching can be described in terms of edge clusters – groups of connections that respond similarly over the course of the scan. Finally, we link these patterns of fluctuations to features present in the movie. Collectively, our findings represent a conceptual bridge between network analysis of time-varying connectivity and naturalistic imaging, and strengthens the link between studies of inter-subject similarity with cognition.

## RESULTS

We analyzed fMRI data from 29 subjects each of which underwent eight scans (four rest and four movie-watching) on two separate sessions, totaling approximately two hours worth of data for every subject. For both scan categories, we estimate time-averaged (static) and time-varying whole-brain FC. Complete methodological details can be found in the section entitled **Materials and Methods.** In this section, we summarize the results of several analyses. First, we compared time-averaged FC between the conditions, identifying differences at the level of individual connections, but also in terms of segregation and integration. Next, we examined time-varying FC and designed a statistical procedure to identify temporal windows during which subjects’ networks become highly similar to each another. We investigated network structure within these windows and discovered that high levels of inter-subject similarity corresponded to decreased modularity and increased dissimilarity with respect to resting FC. Next, we investigate the edge-level correlates of movie-watching, leveraging hypergraph clustering to uncover constellations of connections that follow similar trajectories during movie-watching. Lastly, we demonstrated that fluctuations in time-varying FC were differentially associated with the presence and absence of specific features in the movie, suggesting possible drivers of brain state changes.

### Time-averaged FC during rest and movie-watching conditions

Time-averaged FC is thought to reflect the strength of communication between pairs of brain regions. The magnitude of coupling can be modulated by sensory input, stimulation, or task constraints. In this section we compare rest and movie-watching to assess what effect, if any, movie-watching has on whole-brain patterns of FC.

Fig. 1*a* and Fig. 1*b*, we show the session-, subject-, and time-averaged FC for the resting and movie-watching conditions, respectively. Visually, these two patterns appear highly similar (the correlation of their upper triangle elements confirms as much *r* = 0.69; *p <* 10^−15^). However, when we compute the average change in each connection’s weight, we find systematic differences between the two conditions (Fig. 1*c*; we show a system-averaged version of the same plot in Fig. S1*a*)^1^. To assess the statistical significance of differences, we compared the observed differences to those obtained using a permutation-based null model in which we uniformly and randomly permuted condition labels (within subjects). In general, we found that most connections (*>*60%) exhibited a statistically significant difference between conditions (Fig. 1*d*; false discovery rate fixed at *q* = 0.01; *p*_*adjusted*_ = 0.0062).

**FIG. 1.**
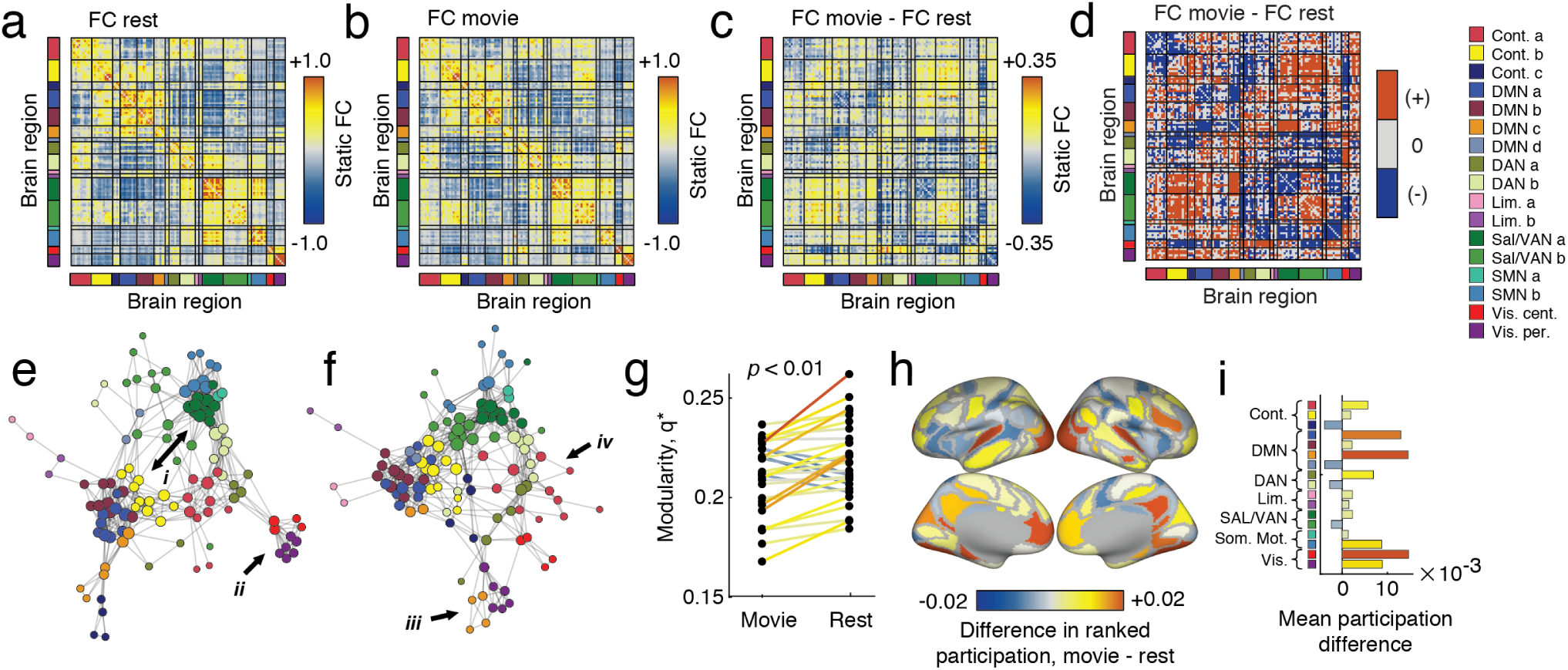
Time-averaged FC at rest *versus* movie-watching. (*a*) Time-averaged FC matrix at rest, ordered by brain systems. (*b*) Time-averaged FC matrix during movie-watching, ordered by brain systems (*c*) Unthresholded mean differences in FC between time-averaged movie-watching and rest. (*d*) Result of statistical analysis comparing movie-watching and rest. Red cells indicate connections that were significantly stronger during movie-watching than rest, whereas blue indicates the opposite. Grey cells were not statistically different between conditions. Panels *e* ad *f* depict thresholded versions of the matrices from *a* and *b*, with node locations determined by a force-directed embedding algorithm. We draw attention to particularly salient differences using Roman numerals. (*g*) Differences in between movie-watching and rest. (*h*) Node-level differences in ranked participation coefficient. (*i*) Same information as in panel *h*, but with differences in participation coefficient averaged by cognitive system.

In general, these differences were distributed across the brain and involved multiple cognitive systems [47]. To help visualize these differences, we generated force-directed embeddings of the task-free and movie-watching networks (Fig. 1*e* and Fig. 1*f*). In both plots, we used arrows to draw attention to particularly salient differences. For instance, in task-free conditions we found that sub-components of the salience and default networks appeared segregated from one another, but that during movie-watching FC between those systems increased so that they were more strongly connected and integrated (Fig. 1*e*,i). Similarly, both sub-components of the visual system appeared highly segregated at rest (Fig. 1*e*,ii), but became more integrated during movie-watching. Other interesting differences included the integration of default mode with visual sub-components (Fig. 1*e*,iii) and the dissolution and distribution of default mode components throughout the network (Fig. 1*e*,iv).

These previous analyses were carried out at the level of individual connections. We also calculated changes in modularity and participation coefficients, measures that index the level of segregation among network modules [48] and the extent to which nodes’ connections are distributed across module boundaries [49], respectively. In agreement with past studies showing that network segregation decreases during tasks [15], we found that modularity, *q\** is greater during rest than movie-watching (Fig. 1*g*). We also found that, as a result of changes in FC, nodes have repositioned themselves with respect to cognitive systems, with areas in the default mode and visual systems becoming more hub-like during movie-watching than at rest (Fig. 1*h* and Fig. 1*i*).

In summary, the findings presented here suggest that movie-watching drives the brain into a more integrated and less modular network architecture. While visual and somatomotor systems are among those whose position within the network changes, differences between the two conditions are distributed and manifest within and across virtually all brain areas and systems. This is an important observation; during movie-watching subjects receive visual and auditory stimuli. One possibility is that differences in network architecture are localized to brain systems associated with the processing of those sensory modalities. However, the involvement of DMN and higher-order association cortices suggest that movie-watching taps into a constellation of brain systems, likely involved in processing more than low-level sensory stimuli.

### Inter-subject similarity and time-varying FC

The previous analysis was carried out on time-averaged FC and revealed differences in network structure over entire scan sessions. To investigate changes over shorter timescales, we calculated time-varying FC using a sliding window technique. Past analyses of time-varying FC networks have focused mostly on properties of time-varying FC defined at the level of individual subjects. Our focus, on the other hand, is on correlated fluctuations in network structure in which subjects’ networks reconfigure in concert with one another. Correlated fluctuations are less likely to originate due to sampling variability and more likely to reflect shared network-level responses to stimuli in the movie [38, 50]. To investigate these kinds of fluctuations, we developed a metric of inter-subject similarity (ISS), which assesses at each time point, the mean similarity of subjects’ FC to one another (Fig. 2).

Because FC tends to be similar across individuals, in general ISS will be nonzero, even in the absence of “true” inter-subject synchrony. To distinguish periods of true inter-subject similarity from baseline, we develop a novel statistical testing procedure. Specifically, we compare mean ISS during movie-watching, when subjects are presented with identical time-locked audiovisual stimuli, with mean ISS during resting conditions, when each subject’s FC fluctuates independently over time. In Fig. 3*a* we show the distribution of ISS values across time for one of the movies. We show an analogous plot for the resting condition in Fig. 3*b* (analogous plots for all four movie and resting scans are shown in Fig. S2). Note that the shape of the distribution fluctuates over time, periodically increasing to reflect windows in time when subjects’ FC becomes highly similar. Rest, on the other hand, is characterized by a distribution of ISS values that changes little over time. To identify periods of high synchrony across subjects, we compared mean ISS at each time point during movie-watching with the distribution of mean ISS values during rest. We note that ISS is related to the concept of inter-subject FC [39] and that, in general, periods of high ISS correspond to periods when inter-subject FC is not equal to zero (Fig. S3).

**FIG. 2.**
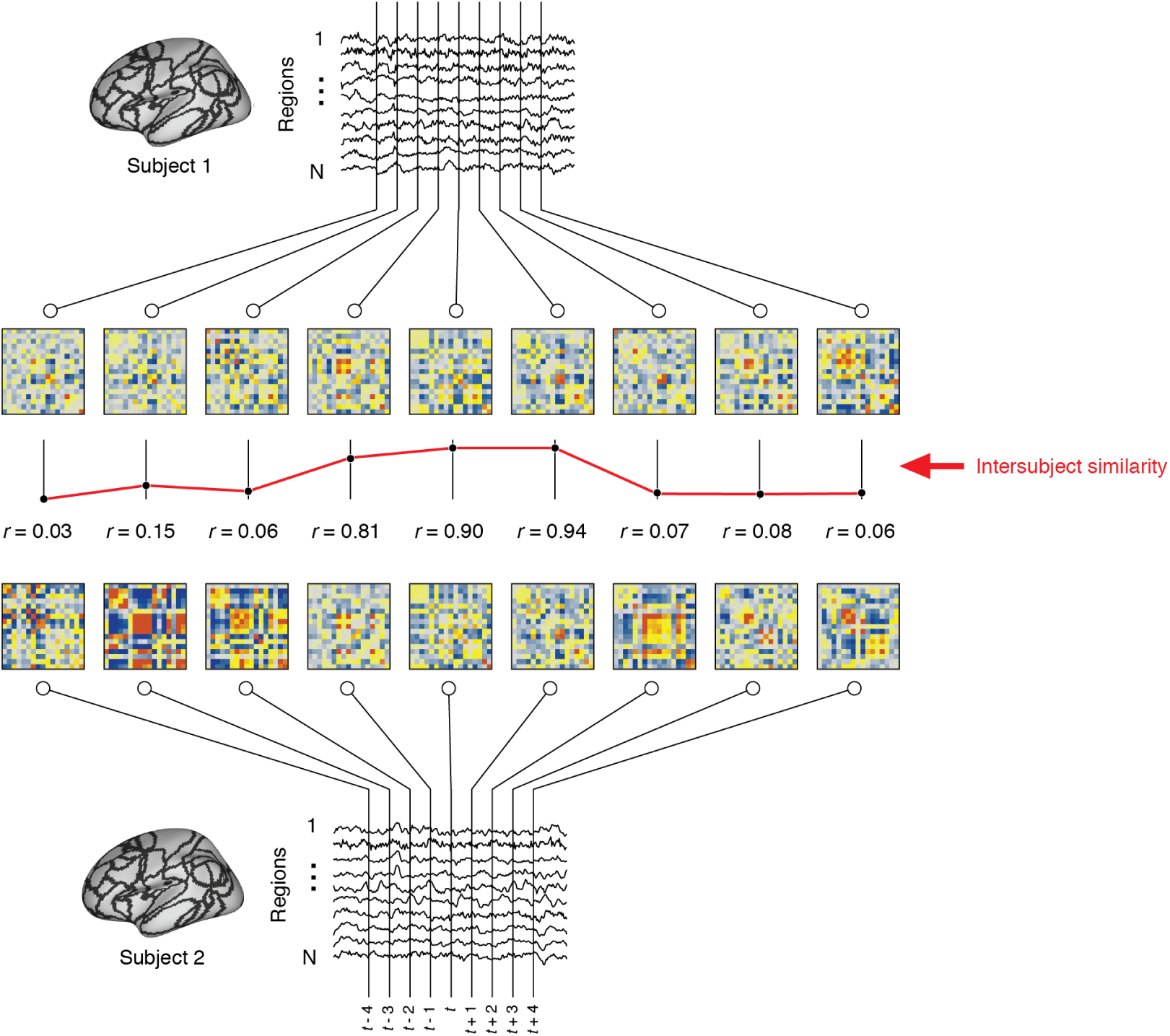
Schematic illustration of inter-subject similarity. We compare network structure between pairs of subject and across time using the inter-subject similarity (ISS) metric. To calculate ISS, we first divide fMRI BOLD time series into overlapping windows and, using only the samples that fall within a window, estimate each window’s FC. Given networks estimated for two subjects at time *t*, we calculate ISS by vectorizing the upper triangle elements of each network and computing the Pearson correlation of those elements. We repeat this procedure for each window in time and for all pairs of subjects. This procedure generates a time-varying estimate of the similarity between subjects’ networks.

**FIG. 3.**
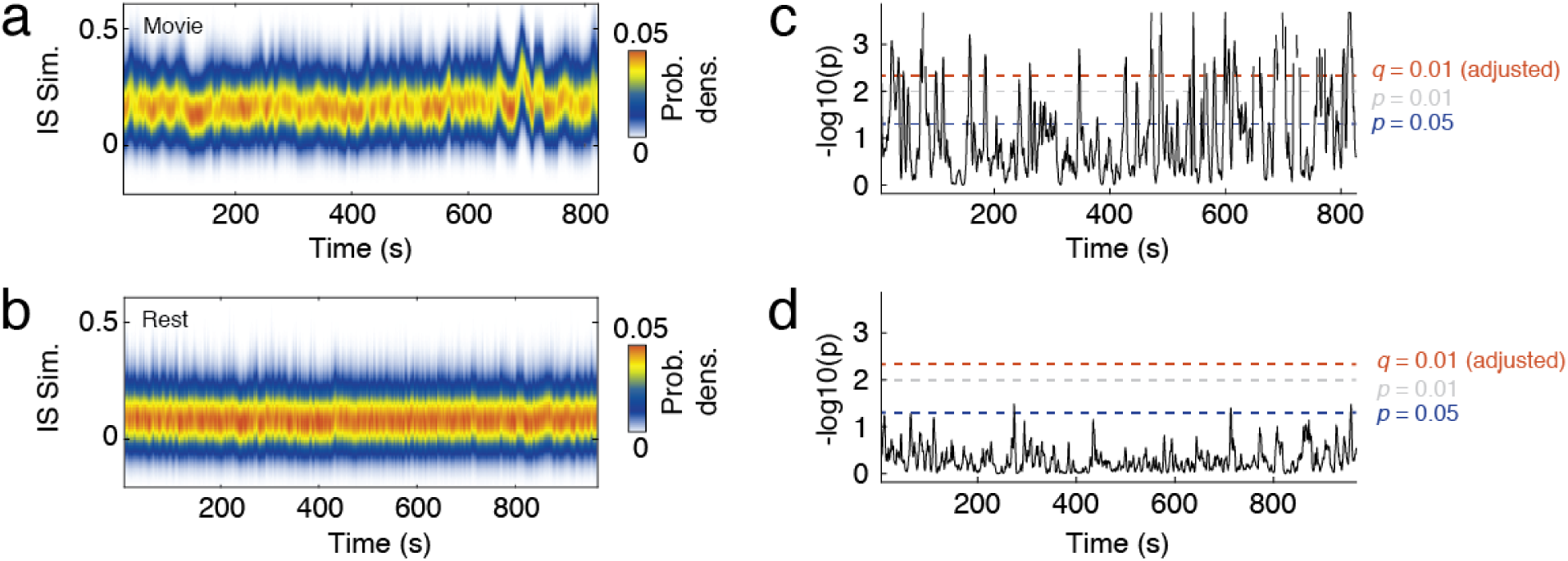
Statistical analysis of inter-subject similarity. (*a*) Distribution of inter-subject similarity (ISS) across time during a representative movie-watching scan. *b*) Distribution of inter-subject similarity (ISS) across time during rest. (*c*) We developed a statistical test to quantify the probability of observing a mean ISS value during movie-watching by chance, where we defined chance based on ISS values at rest averaged across time. This procedure associated each moment in time with a *p*-value, against which we performed statistical testing. The three dotted lines represent different critical values, ranging from *p* = 0.05 (uncorrected for multiple comparisons) to an adjusted *p*-value after fixing false discovery rate at *q* = 0.01 (1%). When we applied these critical values to movie-watching data, we consistently identify periods of time that exceed the criterion for statistical significance. (*d*) Applying the same criteria to resting data (using ISS values from 3/4 scans to estimate *p*-values for the remaining scan), we find no points in time that exceed the critical value.

Using this procedure, we estimated a *p*-value at each time point corresponding to the probability that the observed ISS value would occur during rest. This procedure resulted in a time-series of *p*-values, which we could use to identify periods when ISS during movie-watching was greater than expected by chance. In Fig. 3*c* we show negative log *p*-values for the movie-watching condition. Note that there exist periods when the *p*-values exceed even stringent statistical thresholds. In contrast, Fig. 3*d* shows analogous *p*-values for one of the task-free scans (the null distribution was estimated from the three other task-free scans to avoid circularity). Note that in this case, the *p*-values never exceed the statistical threshold. This observation suggests that, not only does this test distinguish periods of time when subjects exhibited high synchrony during movie-watching, but also demonstrates high specificity by accurately detecting no ISS during rest.

### Inter-subject FC fluctuations are associated with fluctuations in modularity and static resting FC

In the previous section we developed a statistical test for distinguishing periods of high and low ISS. We applied this test to time-varying FC estimated during movie-watching and identified periods of time when ISS was greater than expected by chance (false discovery rate controlled at *q* = 0.01; *p*_*adjusted*_ = 6.3 *×* 10^−4^). On average across all movies, this procedure detected 22.8 *±*14.2 periods of duration 3.6 ± 2.4 seconds (4.5 ± 2.9 TRs)^2^. We note that in-scanner head motion (framewise displacement) was not significantly different between periods of high and low ISS (Fig. S4) and that ISS was not sensitive to global signal regression (Fig. S5).

Periods of high ISS are, by definition, separated by periods of low ISS. What features of brain networks distinguish these periods of time from one another? To address this question, we extracted representative networks for each subject and for every period of low/high ISS. We defined a representative network to be the network

within a given low/high ISS period with the greatest average similarity to other networks within the same window, ensuring that representative networks were all estimated using the same number of samples and not biased by the duration of a given low/high ISS time period.

For each representative network we measured two quantities: first, its modularity, *q*\* [51], a measure of the level of segregation among a network’s communities, and also its similarity to time-averaged FC. In terms of modularity, we found that periods of high ISS were associated with decreased modularity (Fig. 4*a*; *p* = 0.0008), suggesting that cognitive systems become increasingly integrated with one another during periods when high ISS. At the same time, we found that during periods of high ISS were associated with reduced similarity to time-averaged FC (Fig. 4*b,c*).

**FIG. 4.**
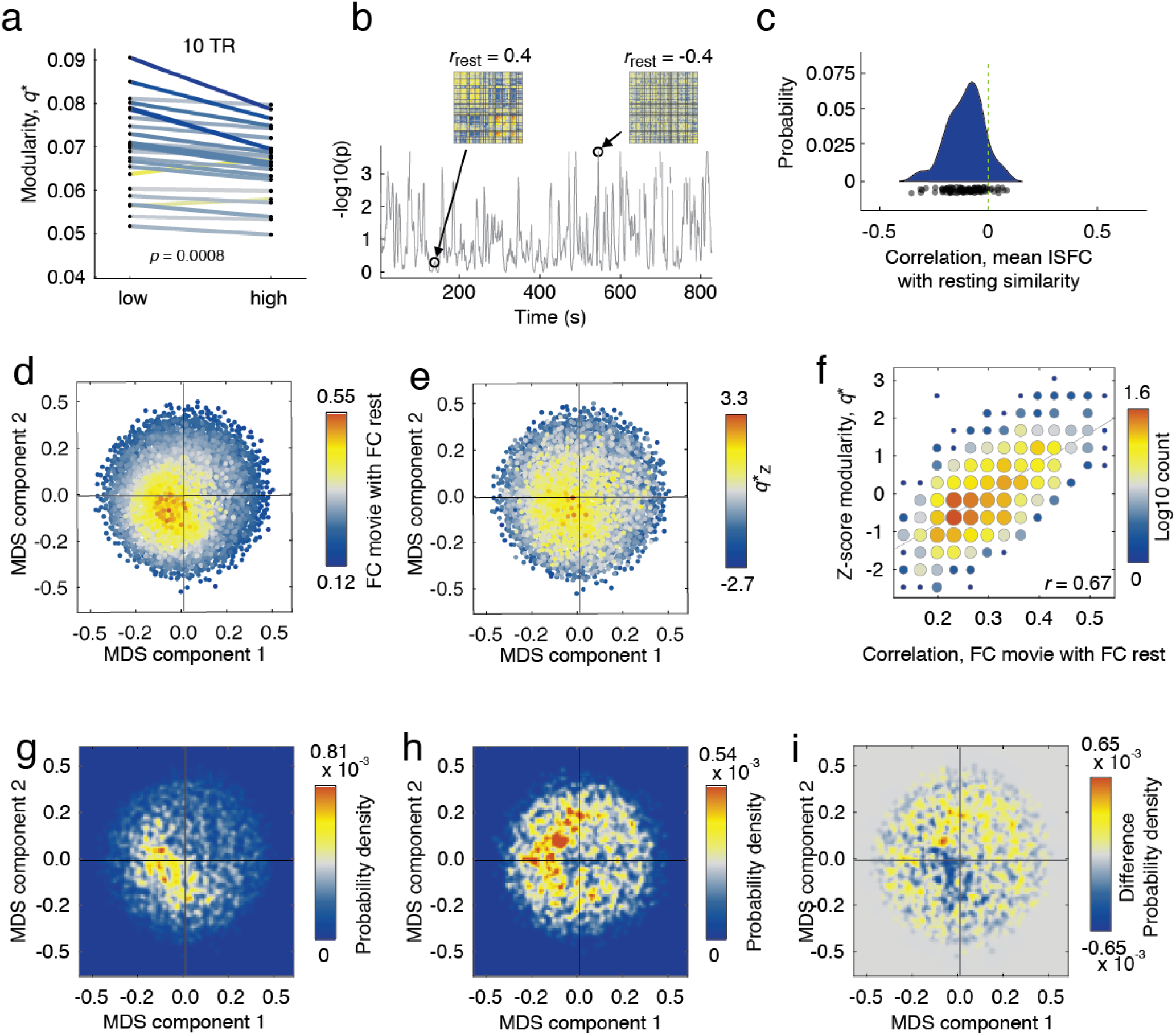
Differences in modular architecture during periods of high and low inter-subject similarity. (*a*) Differences in modularity during low *versus* high inter-subject similarity. (*b*) We calculated the correlation of subjects’ functional connectivity at rest with their time-varying functional connectivity for every instant in time. (*c*) Distribution of correlation coefficients from previous analysis. We used multi-dimensional scaling to embed subjects’ functional connectivity matrices in two dimensions. In panels *d* and *e* we show the same set of points, each corresponding to functional network and colored according to their similarity with resting functional connectivity and modularity. In panel *d*, for instance, brighter points indicate greater similarity to resting FC. Similarly, in panel *e*, brighter points indicate greater levels of modularity. (*f*) Scatterplot of resting correlations with modularity. (*g*) The density of points corresponding to low inter-subject similarity coincides with functional networks that are both modular and correlated with static rest. (*h*) Points corresponding with high inter-subject similarity are distributed along the perimeter. (*i*) Difference in distributions.

To better visualize changes in network organization during high and low ISS, we projected subjects’ FC patterns in two-dimensional space using multidimensional scaling, which reduces the dimensionality of these data while approximately preserving distance structure. In Fig. 4*d* and Fig. 4*e* we show those low-dimension embeddings with points colored according to two different criteria. In Fig. 4*d*, each point is colored according to its similarity with respect to the time-averaged resting FC. In Fig. 4*e*, points are colored according to their modularity (z-scored within subjects to remove subject-specific differences in baseline modularity).

Note that the brightest points in both plots largely coincide with one another, indicating that modular networks are also more rest-like. We further quantified this relationship by computing the correlation of modularity with the similarity to time-averaged, task-free FC and found that the two were strongly related to one another (Fig. 4*f*; *r* = 0.67; *p <* 0.01).

We investigated this relationship further by categorizing each point in Fig. 4*e*,*f* according to whether it corresponded to a period of high or low ISS. We found that periods of low ISS overlapped closely with one another and were concentrated near the middle of the embedding diagram (Fig. 4*g*). Periods of high ISS, on the other hand, were distributed around the border of the embedding (Fig. 4*h*). We show the difference in these density plots in Fig. 4*i*.

In summary, we found that subjects oscillate between periods of high and low similarity, high and low modularity, and network architectures that are rest-like and dissimilar from rest. Surprisingly, all of these changes were correlated with one another. One implication of these findings concerns the link between modularity and cognitive processing. Past studies have reported that when subjects perform cognitively-demanding tasks, their network modularity decreases [15, 16, 18]. Here, we observe that same network-level signature, but because movie-watching is unconstrained, we lack a direct link to cognition. On the other hand, the decreases in modularity took place when network architecture was least similar to rest and subjects were most similar to one another, suggesting that subjects were responding in concert to some aspect of the movie stimulus.

### Movie-watching organizes functional connections into temporal communities

In the previous sections, we showed that movie-watching induces changes in network structure compared to rest and that, when FC is resolved at finer temporal scales, FC oscillates between distinct brain states. It remains unclear, however, which functional connections drive this effect and whether those connections exhibit consistent organization across movies. To address this question, we used a data-driven clustering method [48] to uncover groups of connections that not only exhibit similar temporal trajectories across subjects, but also follow similar trajectories across time [52, 53]. This approach involved calculating subject-averaged time-varying FC, yielding a time series for every edge. We then estimated the edge-by-edge correlation matrix, which we clustered using modularity maximization [54, 55]. This procedure resulted in the detection of twelve large communities comprised of edges that responded similarly across subjects and were also correlated with each other. We show the results of this procedure in Fig. 5.

**FIG. 5.**
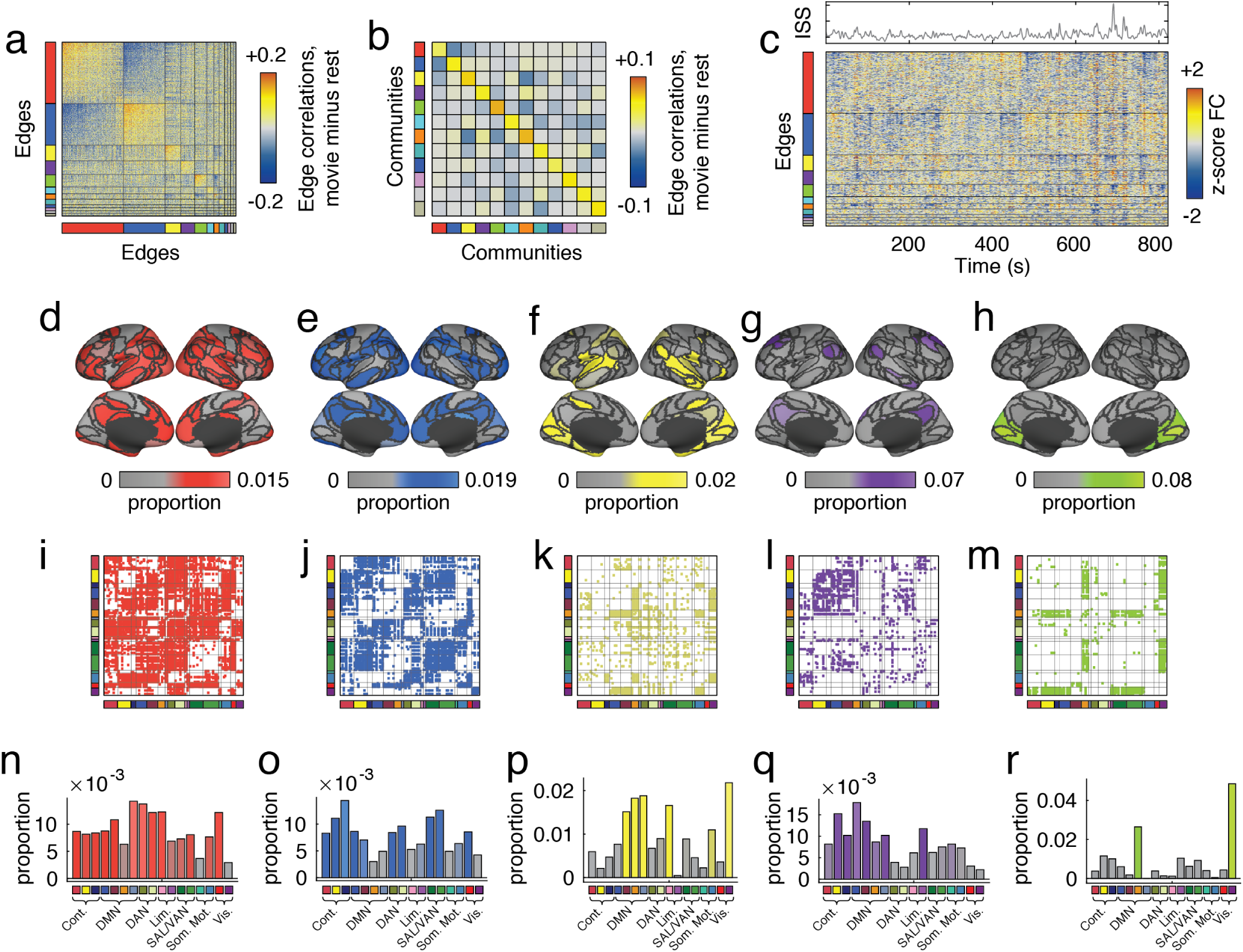
Detecting edge communities. Panels *a* and *b* depict edge correlation matrix and the community-averaged edge correlation matrix. Rows and columns are ordered by community. Here, we exclude small and singleton communities. (*b*) Edge time series organized by community label. To better understand which parts of the brain were associated with each of the five communities, we calculated the fraction of all edges assigned to community that were incident upon each brain region. Panels *d*-*h* show these proportions projected onto cortical surfaces. We also visualize the edge communities by labeling edges in the connectivity matrix by community (panels *i* - *m*) and by aggregating node proportions according to cognitive system (panels *n*-*r*).

Fig. 5*a* depicts the edge correlation matrix that we used as input to the modularity maximization algorithm ordered according to detected communities. Fig. 5*b* depicts the same matrix after averaging connection weights within and between pairs of community. We show the edge trajectories in Fig. 5*c*. Similar trajectories for all movies are shown in Fig. S7.

To further characterize each community, we mapped edges to the nodes upon which they were incident. Because some of a node’s edges can be affiliated with one community and other edges affiliated with another, this means that brain regions can participate in multiple communities simultaneously. We show the five largest edge communities mapped back to the level of nodes and onto brain regions in Fig. 5*d* -*h* (the remaining communities and community overlap indices are shown Fig. S8, Fig. S9, and Fig. S10). The first community was the largest, and was made up of 2175 edges (34% of all possible) and included mostly higher-order association cortex while excluding sub-components of visual and somato-motor systems (Fig. 5*d*). The second community (1473 edges or 23%) involved similar brain areas and systems, but a different collection of edges (Fig. 5*e*). This observation suggests that these higher order association areas are not unifunctional in the context of movie-watching, but play multiple, dissociable roles. Figs. 5*f-h* depict communities three, four, and five. These communities were comprised of 558, 490, and 435 edges (8.7%, 7.6%, and 6.8% of all connections), and involved default mode and visual systems, default mode and limbic, and the visual system, respectively. We present matrix visualizations of temporal communities in Figs. 5*i-m* and, for each community, the proportion of edges associated with canonical brain systems that make up each community (Fig. 5*n-r*).

Collectively, these observations suggest that movie-watching does not induce changes in the weights of single, independent connections. Instead, movie-watching leads to correlated fluctuations among groups of edges. These groups are not simple recapitulations of known cognitive systems, but involve multiple systems, suggesting that movie-watching demands from a fixed neural substrate a multitude of functions. Similar to other studies that reported high levels of variability in somatomotor cortex [56–58], we find that somatomotor cortex is among the systems with greatest participation across communities. This agrees with other studies that have reported activations in motor cortex under diverse contexts, including observing actions of others [59], planning and execution of imagined movements [60], and social cognition [61]. From a methodological perspective, these findings complement other approaches for identifying patterns in brain network organization [7, 8, 11, 47]. Unlike these past efforts, which imposed non-overlapping partitions over brain regions, the approach used here maintains an advantage in that it allows for brain regions to be affiliated with more than one community.

### Linking inter-subject similarity to movie features

In the previous section, we showed that functional connections followed clustered trajectories across time during movie-watching. However, these results do not directly identify the movie features to which the subjects were responding. In this section, we investigate the relationship of features from the movies to time-varying fluctuations in FC. We use a multi-linear modeling approach to identify statistical associations between connection weight and the following hand-coded features: presence of human (in any context), human faces and voices (specifically), inter-personal interactions, mean luminance of projected images, and transitions between films (blank screens) (Fig. 6*a*). This analysis returns a regression coefficient for each edge in the network, indicating the sign and magnitude of correspondence between the feature time-series and edge weight fluctuations.

**FIG. 6.**
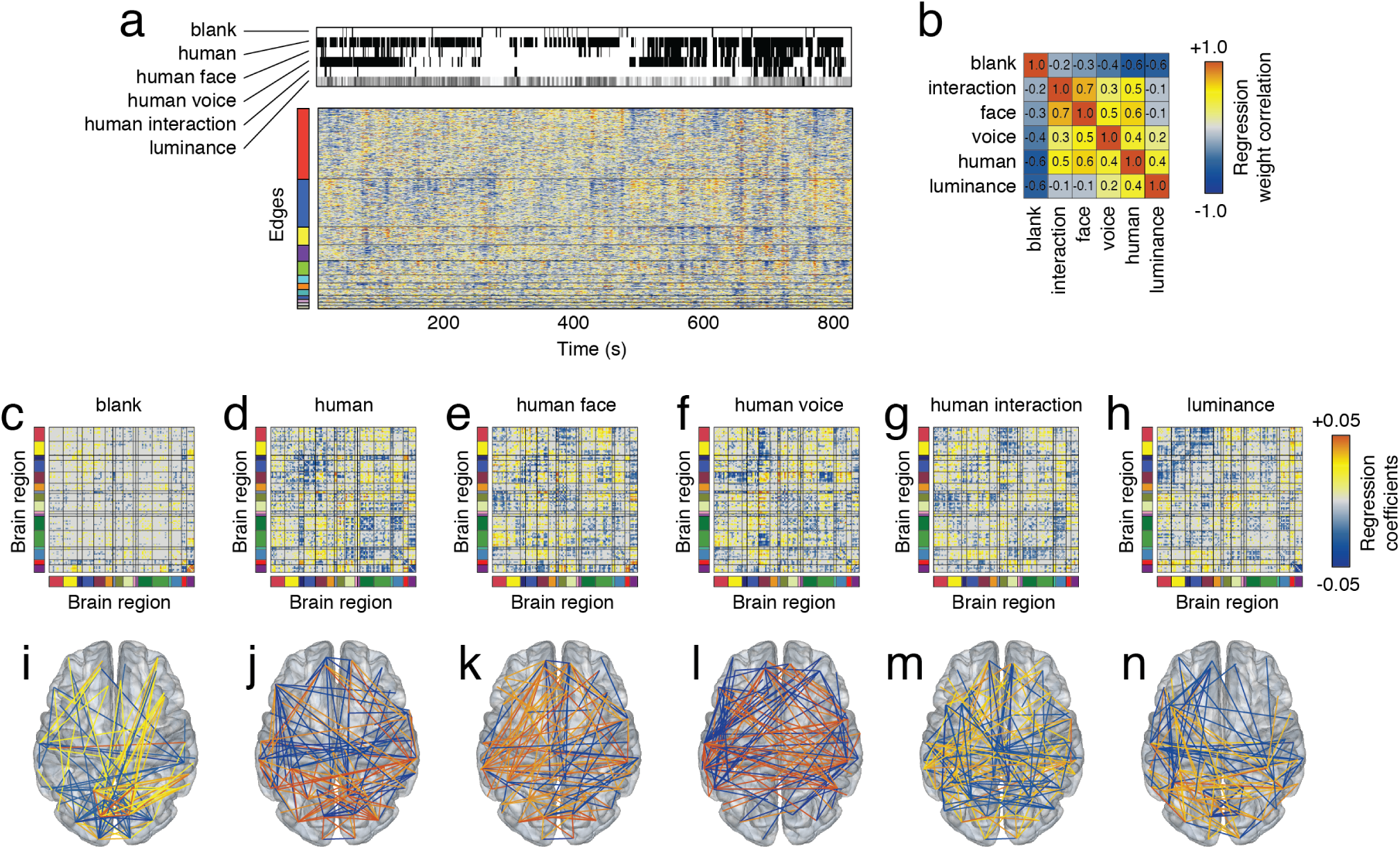
Relating movie features to time-varying CC. We hand-coded and tracked six features across all movies. These included blank screens, presence of a human, a human face, human voice, interactions between humans, and the luminance of the projected image. (*a*) Subject-averaged time-varying FC alongside feature time series. We used linear models to identify combinations of features that predicted subject-averaged time-varying FC. In *b*, we show the correlation of regression weights for each of the six features. (*c* - *h*) Regression coefficients plotted in matrix form. Bright red and yellow elements indicated that the presence of a feature resulted in increased FC, while dark blue indicated that the presence of a feature resulted in decreased FC. (*i* - *n*) The top 2% elements (by absolute value) from each of the six matrices.

In general, we found that each feature was associated with a distinct template of connections (Fig. 6*b*). Most dissimilar was the constellation of edges associated with the black screens that were interspersed between movies, which we label “blank”. Though we found evidence of brain-wide associations, the strongest regression coefficients were concentrated within the peripheral visual system (Fig. 6*c, i*). This is in contrast to the homocentric features (presence of “human”, “human face”, “human voice”, “human interaction”), whose regression weights were all modestly correlated with one another (reflecting, in part, the fact that those regressors were, themselves, correlated; see Fig. S12). These features were associated with spatially-distributed yet feature-specific patterns of connections. For instance, in the case of “human voice”, sub-components of the default mode (DMNb) strongly decouple from the control network and a sub-component of the salience/ventral attention network, but increase connection strength to other default components along with the dorsal attention network (Fig. 6*f, l*). The appearance of human faces, on the other hand, were accompanied by increased coupling between visual and dorsal attention networks and lacked the strong decoupling of DMNb with the salience/ventral attention network. We note that these observations are consistent irrespective of whether our linear models predict FC using each feature independently or whether features are combined into a larger model (see Fig. S11 for comparison of regression coefficients from both models and Fig. S12 for details of how each model deals with correlated regressors).

These observations elaborate on results from the previous section and suggest that not only do groups of connections follow similar trajectories across time, but system-specific sets of connections fluctuate in tandem with the appearance and disappearance of specific features in the movie. This observation, then, serves as a link between time-varying changes in network architecture to features present in the movie.

## DISCUSSION

In this report we aimed to investigate the functional network organization during movie-watching. We found that over long timescales (static FC) and compared to rest, naturalistic stimuli induced increasingly integrated networks and systematic changes in hub structure. Over short timescales, we focused on periods when time-varying network structure appeared highly similar across subjects, which we argue corresponds to periods when subjects may be engaged in similar cognitive processes. We developed a statistical framework for identifying such periods, and applied it to movie-watching and resting state data, where it correctly identified periods of high inter-subject similarity during movie-watching but none during rest. In the case of movie-watching, we found that periods of high inter-subject similarity correspond to highly integrative brain states, in which low intersubject similarity corresponds to network architecture that was increasingly rest-like and more segregated. We also showed that during movie-watching individual functional connections follow similar trajectories and can be clustered accordingly. Finally, we show that time-varying FC coincides with the presence and absence of particular features in the movie, suggesting possible drivers of time-varying FC.

### Naturalistic stimuli as a filter for detecting network events in time-varying FC

Many studies have investigated the time-varying architecture of functional brain networks [19], characterizing the persistence and variability of individual connections [62], uncovering flexible brain regions that change modular assignments across time [56, 63], clustering networks into “states” based on their recurrence structure over one or many scan sessions [57, 64], or relating changes in functional network structure back to anatomical connectivity [25, 65]. Implicitly, these and similar studies have regarded observed time-varying fluctuations in FC as meaningful in some way, reflecting either changes in cognitive state, underlying neurophysiological processes [66], or some combination of the two. However, it remains unclear whether these assumptions are, in fact, warranted. Recent studies have shown that sampling from time-invariant correlation structure produces variability consistent with patterns of observed time-varying FC [31, 32], while the observation that FC is stable both during sleep [67] and under anesthesia raises the question of to what extent an individual’s cognitive state is manifest in FC [68], leading to debate over the verisimilitude of observed time-varying FC [33].

Here, we develop a novel statistical framework to help us better understand which fluctuations are more or less likely to be spurious. To this end, we use a measure of inter-subject similarity as a filter for deciding whether time-varying changes in FC are more or less synchronous across subjects than we would expect. This measure, though it may be conservative and likely fails to identify meaningful but individually variable and idiosyncratic fluctuations in time-varying FC, nonetheless presents a statistical argument that there exist co-fluctuations in time-varying FC not easily explainable were time-varying FC simply reflecting sampling variability from a stationary correlation structure.

These findings have implications for our interpretation of time-varying FC not just during passive movie-watching [35], but also during task-free conditions. Most importantly, we show that the fMRI BOLD response is capable of resolving meaningful fluctuations in time varying FC (albeit only in the statistical sense). This is an important observation, as previous studies have suggested that, combined with the windowing procedure, the slow hemodynamic response is not sufficiently temporally resolved to detect moment-to-moment fluctuations in FC [32]. This suggests that, at least in principle, similar fluctuations should be detectable at rest, which would provide critical validation of studies claiming that some fraction of observed fluctuations in FC at rest are cognitively relevant [26, 29, 30].

### Integrated and segregated brain states during movie-watching

One of the emerging themes in network neuroscience is that the human brain negotiates a careful balance between segregated and integrated states of information processing [69]. This balance is believed to be a critical ingredient for complex, adaptive behavior – a fully integrated system may not be able to support specialized function, whereas a system composed of completely autonomous sub-units may lack the flexibility to perform complex behaviors [70].

This balance gets reflected in the organization of brain networks [7, 16, 17, 71], which can be decomposed into modules that reflect known functional systems. Connections within and between these systems get refined, strengthened, and weakened to support ongoing task demands [15]. These refinements to connections’ weights are far from random, and follow a simple rule that ifluences the extent to which sub-systems are more or less segregated from one another: when an individual is tasked with performing a complex cognitive task, within-system connections weaken while between-system connections become stronger. That is, with increased cognitive load, systems become more integrated and less segregated [72].

Here, we used the modularity metric to assay the extent to which systems are segregated from one another [48], and observed that modularity was closely related to the level of inter-subject similarity. Specifically, when subjects networks were more similar to one another, their networks were less modular (less segregated) compared to periods of time when subjects’ networks appeared dissimilar. This observation is interesting for several reasons. First, it partially corroborates the implicit assumption that network inter-subject similarity indexes periods of shared or similar cognitive processing. We make this claim based on past studies that reported similar decreases in modularity while subjects were explicitly instructed to engage in cognitively complex tasks [15, 16].

Second, this observation forces us to reinterpret other studies that have documented time-varying changes in modularity at rest [22, 23, 62]. In those studies, it was speculated that changes in modular structure across time might be driven by corresponding changes in cognitive state. Validating this hypothesis, however, was impossible due to the unconstrained nature of resting-state. Here, we observe similar patterns during movie-watching, which we *can* partially validate by showing that they occur in synchrony across individuals, suggesting that those fluctuations have similar etiological origins.

In summary, we show that brain network segregation, as measured by the modularity metric, tracks the similarity of network structure between different individuals, a property it shares with task-evoked FC. This observation suggests that periods of high inter-subject similarity may correspond to periods when subjects are attending or responding to stimuli in the movie in similar ways. Nonetheless, we also observed reduced segregation during periods of low inter-subject similarity, whose origins remain unclear. Future experimental work should investigate idiosyncratic components to time-varying FC and concurrent mental operations.

### Temporal segregation through edge clustering

While segregation and integration are usually treated as properties of a network’s nodes, we also studied segregation from the perspective of edges in the network. Specifically, we focused on temporal segregation. That is, we partitioned connections into clusters according to the similarity of their trajectories over time, an approach adopted by several similar studies [52, 53]. Because each node in the network maintains many connections and because clusters were defined at the level of these connections, it was possible for a node to be associated with multiple clusters.

Some of these clusters were largely unimodal and resembled the known system-level architecture of brain networks [8, 10, 11] (see, for example, the visual cluster depicted in Fig. 5h and the somatomotor clusters shown in Fig. S8). On the other hand, we found clusters comprised of multiple brain systems, sometimes dominated by association cortex (for example, the clusters shown in Fig. 5e,g), but other times involving mixtures of association and sensory cortices (the clusters in Fig. 5d,f).

Collectively, these observations suggest that brain network segregation is not simply a static, connectional property, but one that is also encoded across time. This observation has been made before, with many studies findings that the level of segregation in brain networks changes over from moment to moment [22, 23, 62]. These studies, however, have several key limitations. Notably, they have focused on segregation during rest, making it difficult to understand what factors might be responsible for driving fluctuations in segregation. Though these studies have modeled network organization across time, the network structure at each instant was encoded through node-node interactions, implying that the community structure was non-overlapping. Here, we focus on time-varying fluctuations during movie-watching, where changes in network structure are presumably driven by stimuli in the movie, and we model encode network structure through edge-edge interactions. This approach relaxes the definition of what it means for brain regions (or connections) to form modules, allowing brain regions to simultaneously participate in multiple clusters. This overlapping cluster organization agrees with our intuitions that brain regions can play multiple functional roles depending upon context, rather than the sometimes brittle definition of clusters as non-overlapping, which can (falsely) reinforce the notion that there is a one-to-one mapping of regions to function.

Neuroscience aside, overlapping community structure should be investigated further. Recent mathematical results have indicated that the mapping of clusters to edge weights is many to one; the implication is that unless we know the true process by which our network was generated (and its relationship with its cluster structure), we cannot unambiguously claim that the detected clusters are “correct” [73]. This fact motivates the exploration of other approaches for grouping a network into clusters [18, 74]. Future work should focus on detailed comparisons of different clustering methods and their implications for understanding brain function.

### Future directions

Here, we developed a time-varying FC framework for studying movie-watching and other naturalistic stimuli. The primary contribution of this framework is that it enables us to partition time into periods when subjects exhibit similar or dissimilar network architectures. Beyond the analyses presented here, this framework opens up several avenues for future research.

One particularly simple extension involves shifting away from periods of time when subjects are more similar than expected to periods of time when they are more *dis*similar than expected. This extension could be useful for identifying idiosyncratic and individualized responses to stimuli. In other words, rather than focusing on fluctuations in FC that are shared across individuals, focus on periods of time when those fluctuations diverge, which could prove useful for fingerprinting [75].

We used a statistic – inter-subject similarity – that measured, on average, how similar all pairs of subjects were to one another according to their whole-brain patterns of FC. This statistic could be made more useful by, instead of averaging over all subjects, clustering the inter-subject similarity matrix, revealing groups of subjects that may be more internally similar than other groups [76]. This approach could be used for differentiating behavioral phenotypes or revealing sub-structure within a broader disorder [77, 78].

### Limitations

This study also has a number of important limitations. One such limitation is the use of a group parcellation to define nodes in the network. Recent studies have shown that subtle misalignments of subjects to such parcellations can induce biases in FC [79]. Other studies have shown that, with enough data, it is possible to generate more accurate subject-specific parcels that improve parcel homogeneity [80, 81]. Future work should investigate this possibility.

Another limitation concerns the inter-subject similarity measure, itself. In computing this measure, we consider the similarity across *all* possible connections. That is, we calculated similarity based on the whole brain. Future work should could be directed to investigate system- or area-specific similarity [39], which could resolve in greater anatomical detail the drivers of inter-subject synchronization.

Yet another limitation concerns the choice of preprocessing pipeline. In particular, we opted to regress out the global gray matter signal from regional time series. This procedure proven effective in reducing the contribution of in-scanner head motion and physiological noise on FC [82, 83] and for centering connection weights [84]. However, the global signal has also been linked to neurophysiological processes [85] and also serves as an index of arousal [66]. This suggests that, to whatever extent the global signal contributes to synchronous fluctuations in time-varying FC across subjects, our analysis may fail to characterize those fluctuations. We note, however, that the precise contribution of the global signal to FC (and the verisimilitude of those contributions) remains an active and highly debated area of research [86, 87] with no clear resolution [88–90]. Future work on this topic, including comparisons of the fMRI BOLD signal with other imaging modalities [91], will help clarify this debate and lead to more refined processing pipelines for estimating both time-invariant and time-varying FC.

## MATERIALS AND METHODS

### Demographics

We analyzed MRI data collected from *N*_*s*_ = 29 subjects (5 female, 24 male; 25 were right-handed). This cohort was male-dominant, as subjects were intended to serve as controls for a study in autism spectrum disorder, which is more common in men than women. At the time of their first scan, the average subject age was 24.9± 4.7 years.

### MRI acquisition and processing

MRI images were acquired using a 3T whole-body MRI system (Magnetom Tim Trio, Siemens Medical Solutions, Natick, MA) with a 32-channel head receive array. Both raw and prescan-normalized images were acquired; raw images were used at all preprocessing stages and in all analyses unless specifically noted. During functional scans, T2*-weighted multiband echo planar imaging (EPI) data were acquired using the following parameters: TR/TE = 813/28 ms; 1200 vol; flip angle = 60*°*; 3.4 mm isotropic voxels; 42 slices acquired with interleaved order covering the whole brain; multi-band acceleration factor of 3. Preceding the first functional scan, gradient-echo EPI images were acquired in opposite phase-encoding directions (10 images each with P-A and A-P phase encoding) with identical geometry to the EPI data (TR/TE = 1175/39.2 ms, flip angle = 60*°*) to be used to generate a fieldmap to correct EPI distortions, similar to the approach used by the Human Connectome Project [92]. High-resolution T1-weighted images of the whole brain (MPRAGE, 0.7 mm isotropic voxel size; TR/TE/TI = 2499/2.3/1000 ms) were acquired as anatomical references.

All functional data were processed according to an in-house pipeline using FEAT (v6.00) and MELODIC (v3.14) within FSL (v. 5.0.9; FMRIB’s Software Library, www.fmrib.ox.ac.uk/fsl), Advanced Normalization Tools (ANTs; v2.1.0) [93], and Matlab R2014b. This pipeline was identical to the **GLM + MGTR** procedure described in [83].

In more detail, individual anatomical images were bias-corrected and skull-stripped using ANTs, and segmented into gray matter, white matter, and CSF partial volume estimates using FSL FAST. A midspace template was constructed using ANTs’ *buildtemplateparallel* and subsequently skull-stripped. Composite (affine and diffeomorphic) transforms warping each individual anatomical image to this midspace template, and warping the midspace template to the Montreal Neurological Institute MNI152 1mm reference template, were obtained using ANTs.

For each functional run, the first five volumes (≈ 4 seconds) were discarded to minimize magnetization equilibration effects. Framewise displacement traces for this raw (trimmed) data were computed using *fsl motion outliers*. Following [83, 94], we performed FIX followed by mean cortical signal regression. This procedure included rigid-body motion correction, fieldmap-based geometric distortion correction, and non-brain removal (but not slice-timing correction due to fast TR [92]). Preprocessing included weak highpass temporal filtering (*>*2000 s FWHM) to remove slow drifts [92] and no spatial smoothing. Off-resonance geometric distortions in EPI data were corrected using a fieldmap derived from two gradient-echo EPI images collected in opposite phase-encoding directions (posterior-anterior and anterior-posterior) using FSL topup.

We then used FSL-FIX [95] to regress out independent components classified as noise using a classifier trained on independent but similar data and validated on hand-classified functional runs. The residuals were regarded as “cleaned” data. Finally, we regressed out the mean cortical signal (mean BOLD signal across gray matter partial volume estimate obtained from FSL FAST). All analyses were carried out on these data, which were registered to subjects’ skull-stripped T1-weighted anatomical imaging using Boundary-Based Registration (BBR) with *epi reg* within FSL. Subjects’ functional images were then transformed to the MNI152 reference in a single step, using ANTS to apply a concatenation of the affine transformation matrix with the composite (affine + diffeomorphic) transforms between a subject’s anatomical image, the midspace template, and the MNI152 reference. Prior to network analysis, we extracted mean regional time series from regions of interest defined as sub-divisions of the 17-system parcellation reported in [11] and used previously [78, 96, 97].

### Naturalistic stimuli

All movies were obtained from Vimeo (https://vimeo.com). They were selected based on multiple criteria. First, to ensure that movies represented novel stimuli, we excluded any movie that had a wide theatrical release. Secondly, we excluded movies with potentially objectionable content including nudity, swearing, drug use, etc. Lastly, we excluded movies with intentionally startling events that could lead to excessive in-scanner movement.

Each movie lasted between 45 and 285 seconds (approximately 1 to 5 minutes). Each movie scan comprised between four and six movies with genres that included documentaries, dramas, comedies, sports, mystery, and adventure. See Table. S1 for more details.

### Connectivity measures

#### Time-averaged FC

Let **x**_*i*_ = [*x*_*i*_(1), *…, x*_*i*_(*T*)] be the vector of activity for region *i*. We calculated time-averaged FC as the Pearson correlation of activity recorded in region *i* and *j*:

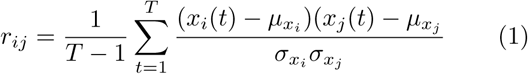

Where 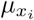 and 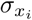 are the time-averaged mean and standard deviation of activity in region *i*. All correlation coefficients were subsequently Fisher transformed.

#### Time-varying FC

In addition to time-averaged FC, we also calculated time-varying FC using a sliding window approach. This involves defining a window length of *L* (in units of TRs) and computing FC using samples within that window only. The time-varying FC at time *t* was calculated as:

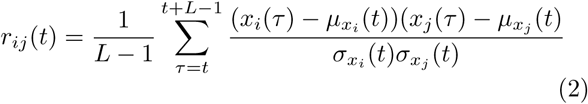

where:

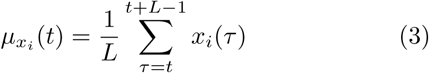

and

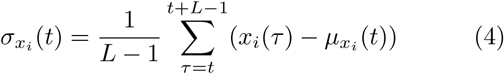

are the mean and standard deviation of region *i*’s activity recording in the *L*-length window starting at time *t*.

An important decision to make in computing time-varying FC using sliding windows is the choice of window length, *L*. Here, we set *L* = 10 TRs (for a window duration 8.13 s). We note that this length is considerably shorter than the durations suggested elsewhere, which have argued that the minimum window duration should be inversely proportional to the slowest frequency component of the fMRI signal. For instance, if filtered to 0.01 - 0.1 Hz, then the shortest window should be 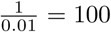 seconds [98, 99]. Our rationale for disregarding this rule of thumb is that the intersubject similarity measure (see the next section) effectively acts as a filter to reduce the rate of false positives. That is, if fluctuations in time-varying FC within a subject are spurious, it is unlikely that similarly spurious fluctuations in another subject occur at precisely the same instants in time.

### Intersubject similarity

Let *r*^*u*^(*t*) and *r*^*v*^(*t*) be the network structure estimated for subjects *u* and *v* at time *t*. We calculate the similarity of these two networks by vectorizing their upper triangle elements and computing the correlation of those vectors: *r*_*u,v*_(*t*). In practice, we compute this measurement for all pairs of subjects and for every window, generating a distribution of intersubject similarity scores that evolves over time.

### Modularity maximization

Modularity maximization is a heuristic for detecting communities in networks [48]. Intuitively, it attempts to decompose a network into non-overlapping sub-networks such that the observed density of connections within sub-networks maximally exceeds what would be expected by chance, where chance is determined by the user. The actual process of detecting communities is accomplished by choosing community assignments that maximize a modularity quality function, *Q*, defined as:

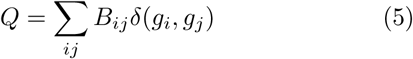

where *B*_*ij*_ = *A*_*ij*_ − *P*_*ij*_ is the {*i, j*} element of the modularity matrix, which represents the observed weight of the connection between nodes *i* and *j* minus the expected weight. The variable *g*_*i*_ is the community assignment of node *i* and *δ*(*x, y*) is the Kronecker delta function, whose value is 1 when *g*_*i*_ = *g*_*j*_ and 0 otherwise. The modularity, *Q*, is effectively a sum over all edges that fall within communities and is optimized when the the observed weights of connections is maximally greater than the expected. In general, larger values of *Q* are thought to reflect superior community partitions.

#### Signed and correlation matrices

In this manuscript, we use two variations of modularity. First, we use the modularity quality function as a means of assessing the level of segregation in a network. Specifically, we estimate a variant of modularity, *Q*\*, which has been shown to be especially well-suited for use with correlation matrices [51]:

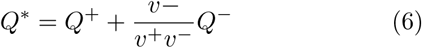

where 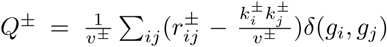. In this expression, 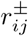 represents either the positive or negative elements of the correlation matrix, 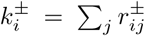, and 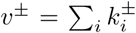. In all instances where we report *Q*\*, it is with respect to the seventeen systems reported in [47].

#### Edge communities

We also apply a second variant of modularity maximization to edge correlation matrices. As a product of calculating time-varying FC, we obtain a time series for each connection in the network, i.e. **r**_*ij*_ = [*r*_*ij*_(*t* = 1), …, *r*_*ij*_(*t* = *T* − *L* + 1)]. From these edge-level time series, we can calculate the edge-by-edge correlation matrix, Ω, whose element Ω{*i, j*}, {*k, l*} is equal to the correlation of **r**_*ij*_ and **r**_*kl*_.

Here, we generate a modularity matrix wherein we treat the edge correlations from the movie as our observed network and the edge correlations from rest as our expected or chance network. This matrix is calculated as: *B*(*γ*) = [Ω^*movie*^ − Ω^*rest*^] − *γ*, where *γ* is a structural resolution parameter that can be tuned to detect different numbers and sizes of communities [100]. We can then optimize the corresponding modularity to identify edges whose trajectories across time are more correlated during movie-watching than during rest. Because this procedure assigns edges to communities rather than nodes, it is possible for a fraction of a nodes’ edges to be associated with multiple distinct communities, so that each node has an overlapping community structure. We note that this approach is similar to the hypergraph clustering reported in [52, 53].

An important open question concerns choosing the optimal value of *γ*. Here, we tested 51 different *γ* values, linearly-spaced between from −0.005 - 0.36. We focused on the value that maximized the minimum average within-community correlation minus the maximum average between-community correlation.

We used a generalization of the popular Louvain algorithm to optimize *Q* [101, 102]. This algorithm is non-deterministic, meaning that different initializations lead to slightly different estimates of community structure (we run the algorithm 100 times). To resolve variability in these estimates we use consensus clustering [103]. The variant of consensus clustering used here is slightly different from that of [103]. Specifically, we calculate a co-assignment matrix from the 100 estimated partitions, whose elements indicate the fraction of times that pairs of nodes were assigned to the same community. In [103], this matrix is iteratively thresholded and clustered until convergence. Here, we estimate the probability that two nodes would be assigned to the same community were community labels randomly and uniformly permuted. Then, we construct a new modularity matrix by subtracting this probability from the co-assignment matrix, which we then cluster using modularity maximization [62, 104, 105].

#### Community overlap entropy

The edge community detection procedure assigned every pair of nodes, {*i, j*} to a community. To better understand how these edge-level labels were related to individual brain areas (nodes), we calculated each node’s community overlap entropy. Let Γ_*i*_ = {*g*_*i*1_,…, *g*_*iN*_} be the set of community assignments for all edges involves node *i*. Each element, *g*_*ij*_ ∈ {1, …, *K*} indicates to which of the *K* communities edge {*i, j*} was assigned. We then calculate the fraction of all edges assigned to each community, *c*, as *Pr*_*c*_, and subsequently calculate the entropy over this distribution as: 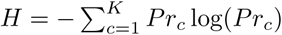. We normalize this entropy to the interval [0, 1] as 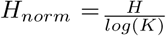. We repeat this procedure separately for every node *i* ∈ {1, …, *N*}, resulting in node-defined entropies, *H*_*i*_.

#### Participation coefficient

Knowing a network’s community structure allows us to classify nodes based on their functional roles. One popular measure for doing so is the so-called *participation coefficient* [49], which measures the extent to which a node’s connections are distributed within or across modules. The participation coefficient is calculated as:

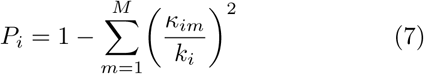

where *k*_*i*_ and *κ*_*im*_ are the total number (or weight) of connections made by node *i* overall and to module *m*.

### Multi-linear models

To map the association of time-varying FC with features present in the movie, we build edge-level multi-linear regression models. Let **r**_*ij*_ = [*r*_*ij*_(1), …, *r*_*ij*_(*T*)] denote the time series of connection weights between nodes *i* and *j*, and let **f**_*l*_ = [*f*_*l*_(1), …, *f*_*l*_(*T*)] be the z-scored time series of feature *l*. All features were coded as binary variables; the exception was luminance, which was coded continuously.

To model the response of connection {*i, j*} to feature, *l*, we constructed the simple linear model:

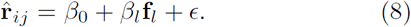

In this equation, 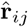 is the predicted connection time series, *β*_0_ is the intercept term, and *β*_*l*_ is the regression coefficient for feature *l*.

We extended this simple linear regression to include multiple terms – one for each of the six coded features:

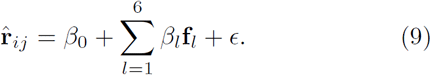

We fit the model to time series data from every connection by minimizing the least square error. The *p*-value associated with each *β* regression coefficient was calculated using a one-sample *t* -test.

## AUTHOR CONTRIBUTIONS

All authors conceived of the study. LB and DPK designed experimental protocols, collected, and preprocessed all fMRI data. RFB and FZE performed all analyses. All authors wrote and approved of the manuscript.

## CODE AVAILABILITY

All data and code are available from the authors upon reasonable request.

## ACKNOWLEDGMENTS

RFB and FZE acknowledge support from Indiana University Office of the Vice President for Research Emerging Area of Research Initiative, Learning: Brains, Machines and Children. This work was supported by the NIH (R01MH110630 and R00MH094409 to DPK and T32HD007475 Postdoctoral Traineeship to LB). For supercomputing resources, this work was supported in part by Lilly Endowment, Inc., through its support for the Indiana University Pervasive Technology Institute, and in part by the Indiana METACyt Initiative. The Indiana METACyt Initiative at IU was also supported in part by Lilly Endowment, Inc. We thank Hu Cheng for MRI protocol development, Soyoung Park for training the FIX classifier, and Brad Caron and Susannah Burkholder for data collection.

---

**FIG. S1.**
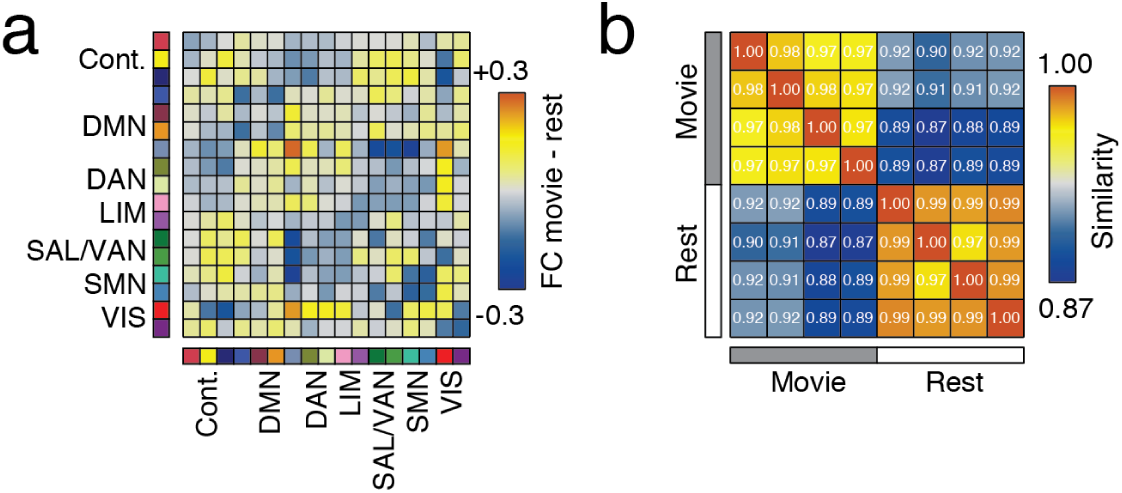
Comparison of time-averaged FC. (*a*) Difference in system- and time-averaged FC between movie and resting conditions. (*b*) Mean similarity of time-averaged movie and resting FC.

**FIG. S2.**
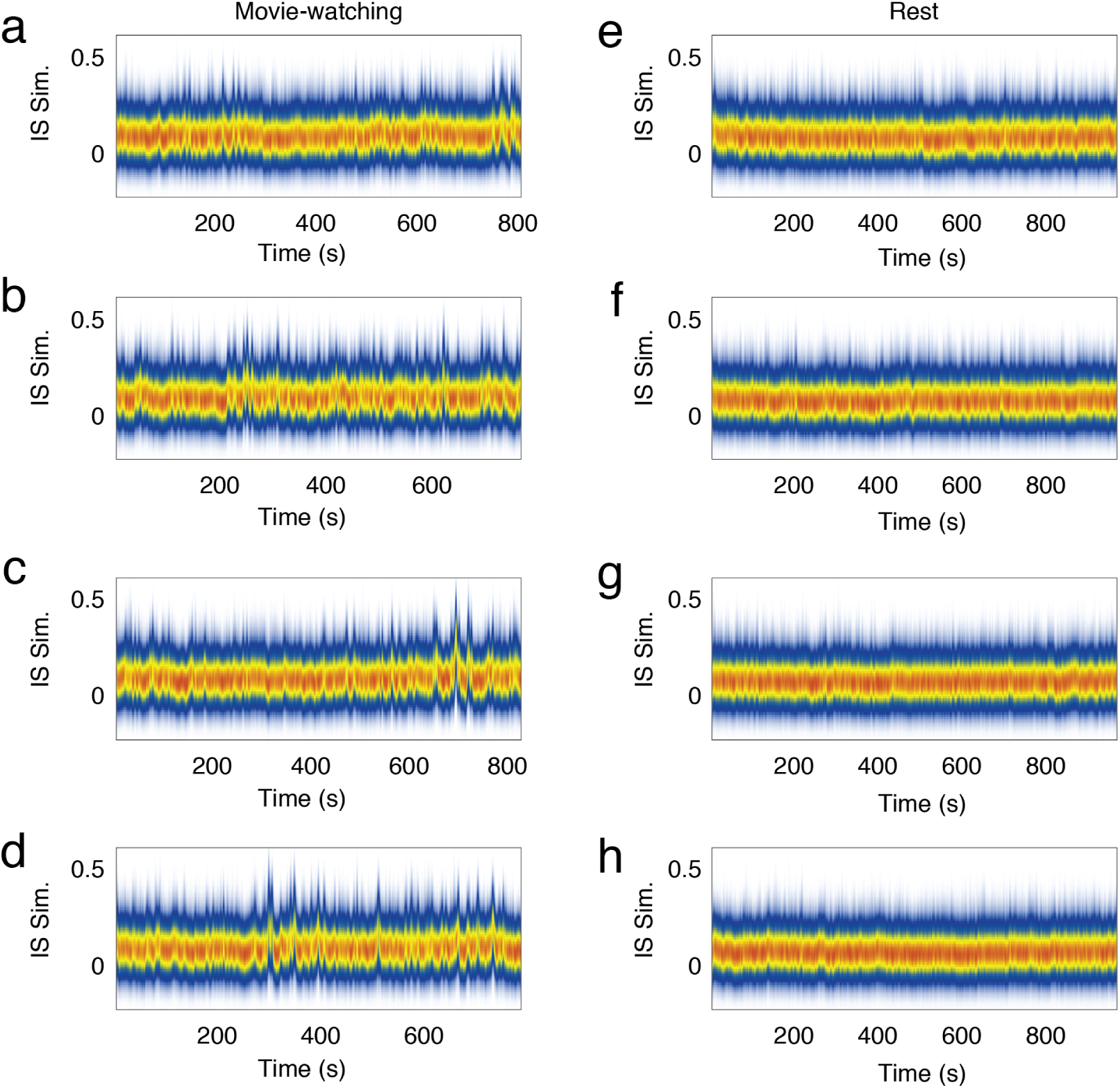
Inter-subject similarity distributions. Panels *a*-*d* depict ISS distributions across time for all four movie-watching scans. Panels *e*-*h*, on the other hand, depict ISS distributions at rest.

**FIG. S3.**
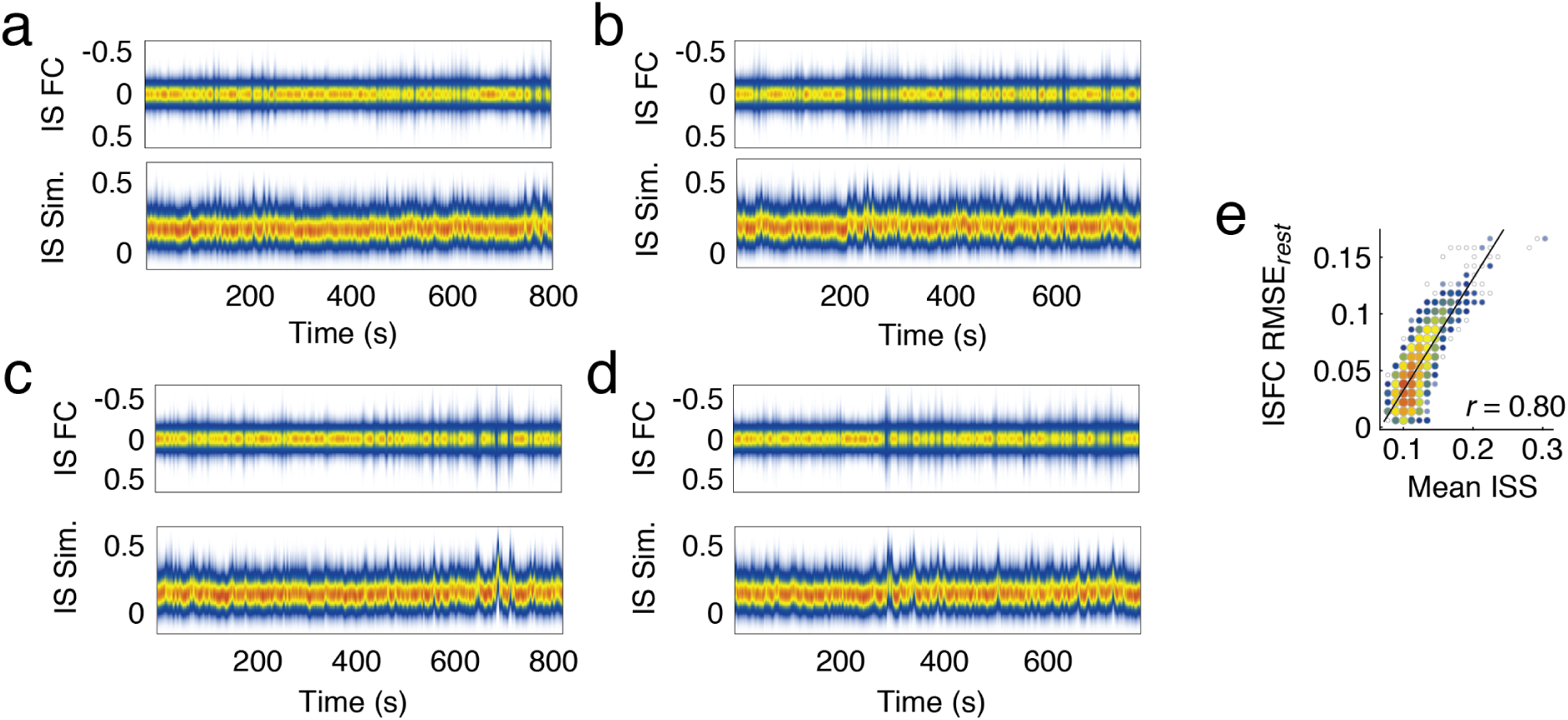
Comparison of inter-subject similarity and inter-subject FC. In the main text, we focus on the measure of inter-subject similarity (ISS). It involves estimating time-varying networks for every individual assessing their similarity at each time point. This approach enables us to identify points in time when network architecture is share across individuals while simultaneously modeling every individual’s whole-brain network, which further allows us to estimate measures like network modularity. An alternative approach for identifying shared network structure involves estimating time-varying inter-subject FC (ISFC), or the correlation of activity in region *i* in subject *s* with region *j* in subject *t* and repeating this procedure for all pairs of regions, all pairs of subjects, and at every time point [39]. Here, in panels *a* - *d*, we show distributions of ISFC alongside ISS. We note that, in general, periods of high ISS coincide with periods of strong (positive or negative) ISFC. In panel *e* we quantify this relationship by computing the correlation between mean ISS and the difference between the ISFC distribution during movie watching compared to rest (root mean squared-error). This indicates that ISS and ISFC, while they relate subjects’ networks to one another differently, identify similarity across subjects at roughly the same points in time.

**FIG. S4.**
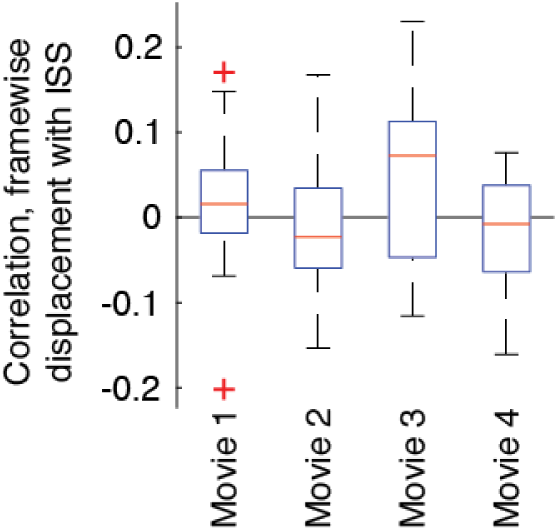
Correlation of in-scanner head motion with inter-subject similarity. We calculated the mean framewise displacement (FD) for each subject within each window (10 TRs or approximately 8.13 s). To assess whether motion might be related to ISS, we calculated the Pearson correlation of each subject’s FD with mean inter-subject similarity (ISS). In general, we found that correlations were consistently centered around to zero for all four movies, suggesting that ISS is not obviously driven by in-scanner motion.

**FIG. S5.**
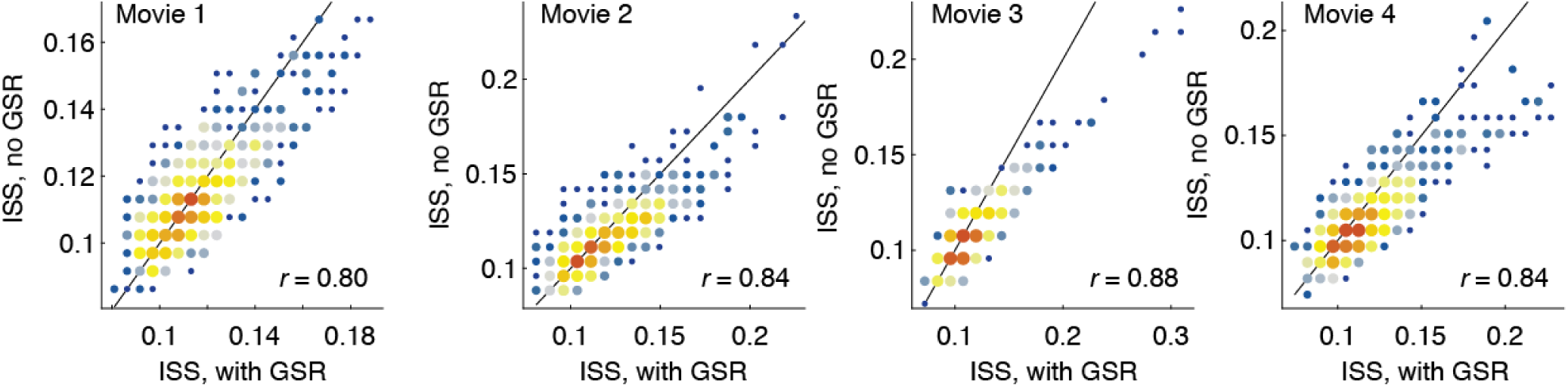
Effect of global signal regression on mean inter-subject similarity for the four movie scans. We calculated inter-subject similarity using data that was processed identically to the what was described in the main text. The only processing step that was omit was the regression of mean gray matter BOLD signal from the data. Here, we compare those ISS measures (labeled ISS, no GSR) wit the ISS measures from the main text (ISS, with GSR). We find that the two are highly correlated, suggesting global signal regression has little effect on ISS.

**FIG. S6.**
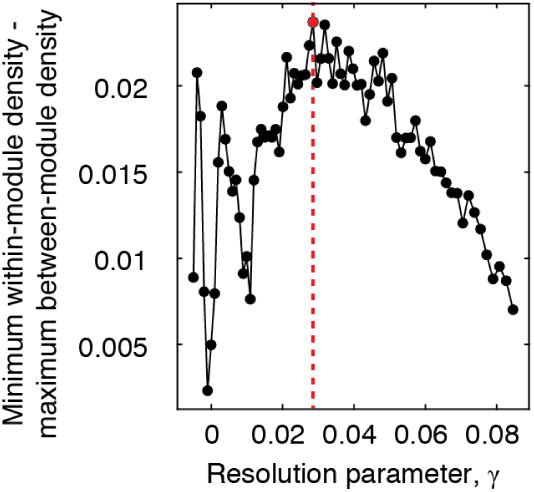
Choice of optimal resolution parameter for edge clustering. The community detection algorithm used to partition edges into communities depends on a resolution parameter, *γ*, whose value determines the size/number of detected communities. We selected *γ* such that the partitions detected at that value produced communities that were maximally segregation. That is, when the minimum internal density of connections across all communities minus the maximum density of connections across communities achieved its peak.

**FIG. S7.**
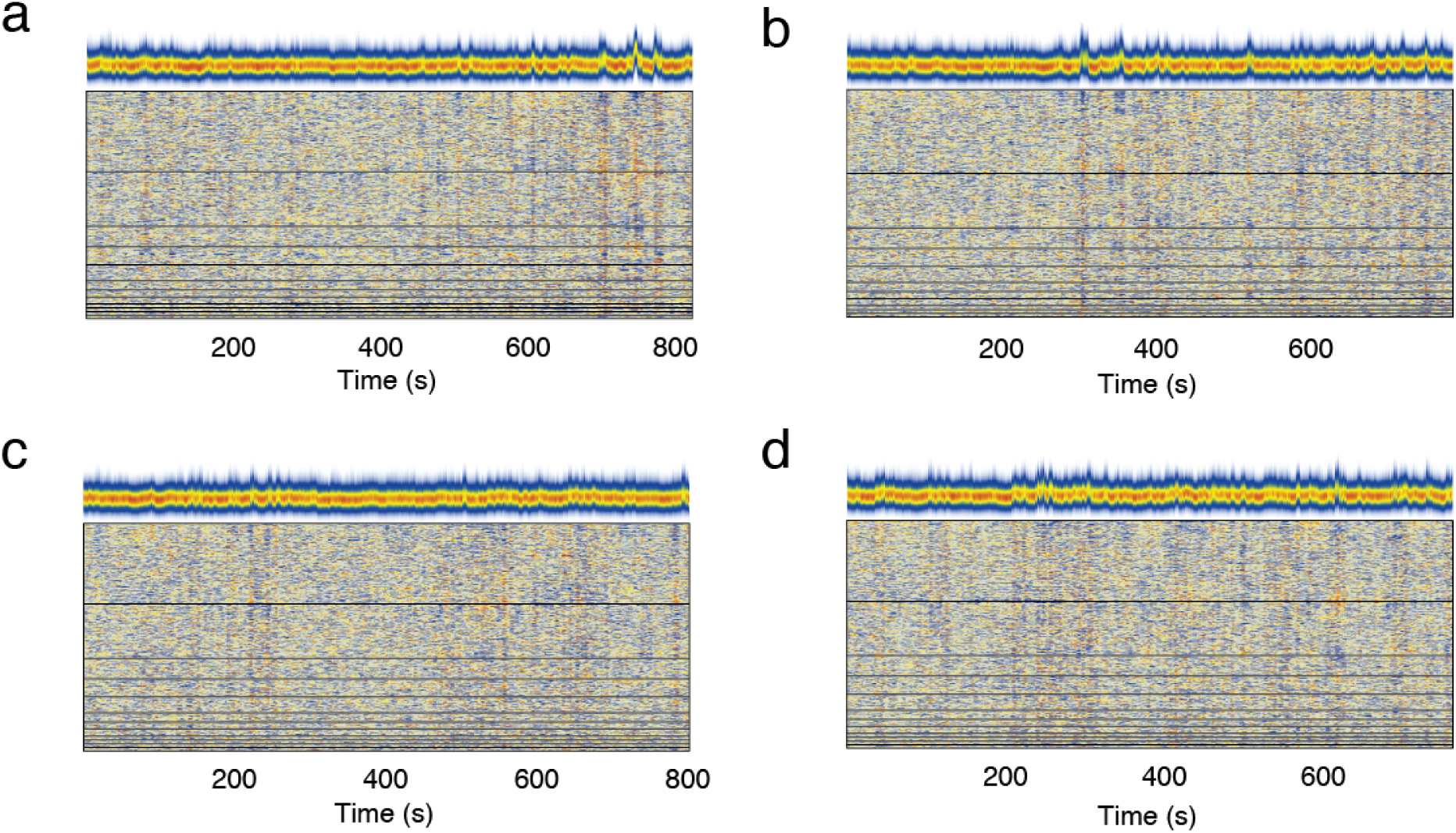
Edge trajectories ordered by community. Panels *a*-*d* depict edge trajectories ordered by community for all four movies.

**FIG. S8.**
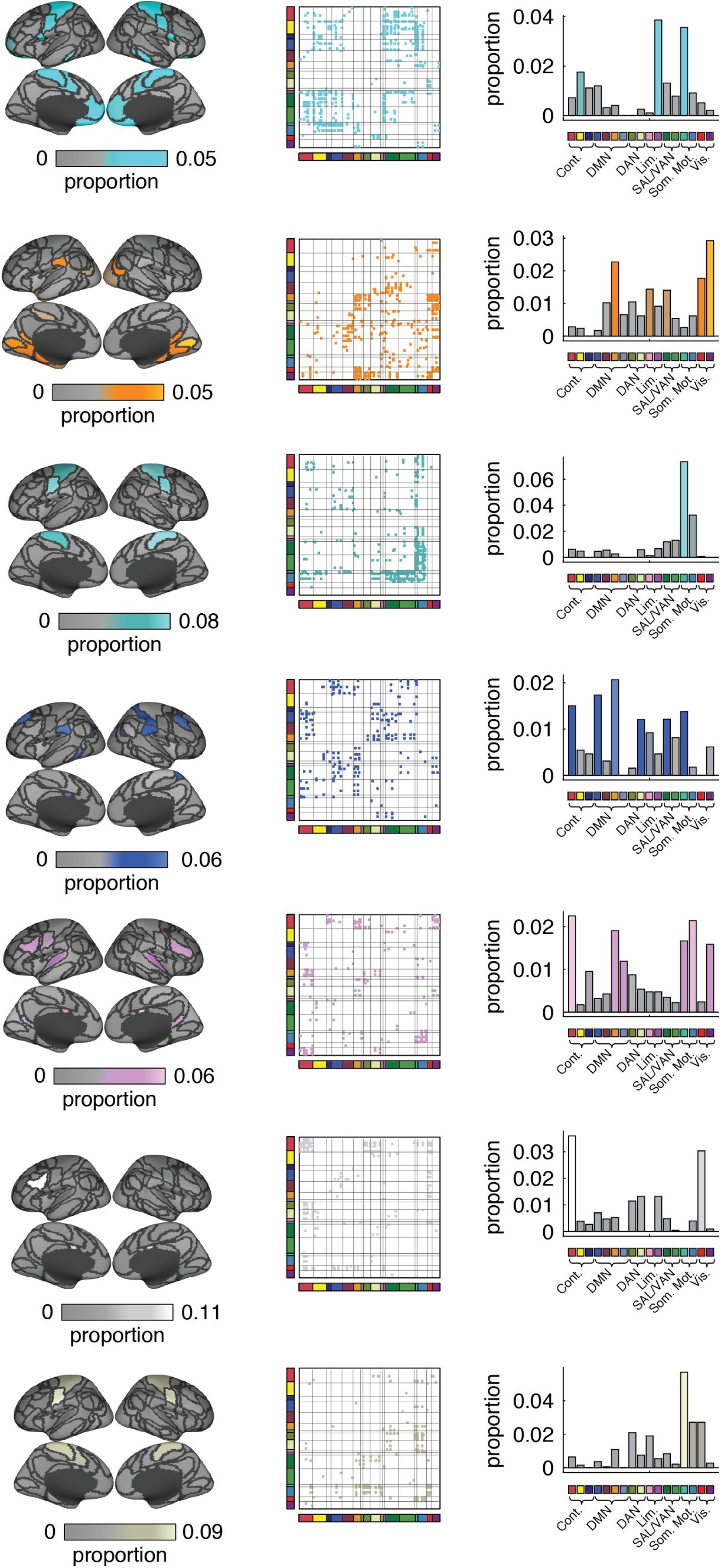
Remaining edge communities. In the main text we showed the five largest edge communities. We show the remaining seven here.

**FIG. S9.**
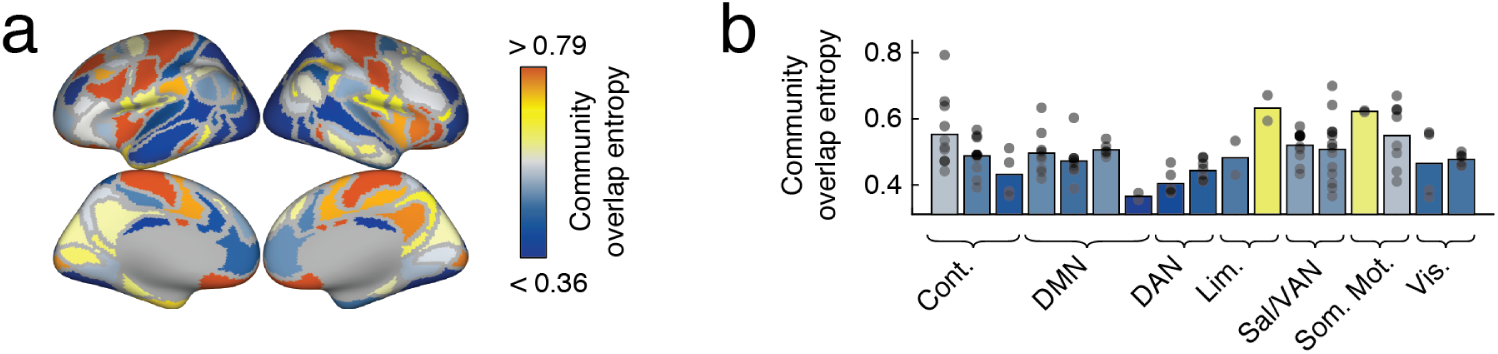
Edge community overlap. For each node we calculated the entropy of its edges’ community assignments. Small values indicate that those edges tended to be assigned to a small subset of communities, while larger values indicate a broader distribution of community labels. In panel *a* we show the entropy values distributed over cortex, while in panel *b* we show those same values aggregated by system.

**FIG. S10.**
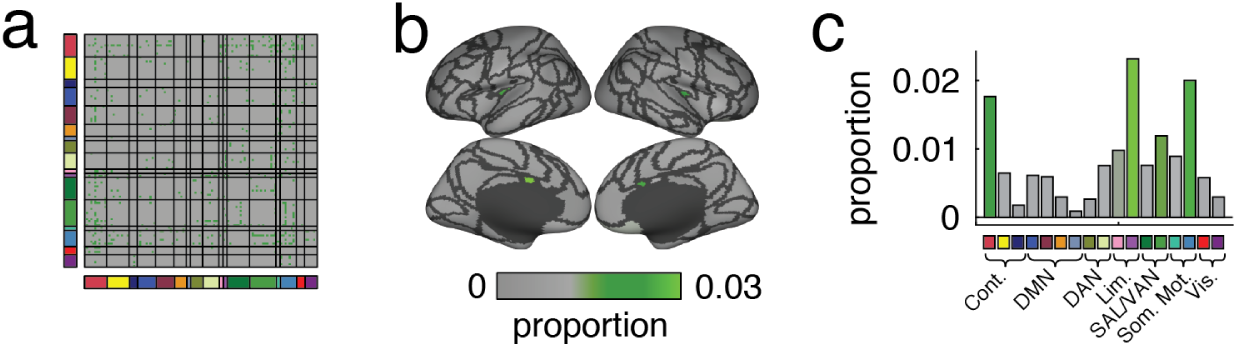
Small and singleton communities. In the main text we clustered edge correlation matrices so that every edge was associated with a community label. We reported the twelve largest communities, however there were many small communities comprised of only a few edges. Here, we aggregate those communities into a single label and provide a short summary of their properties. (*a*) Edges assigned to these communities. (*b*) Projection of those connections onto brain areas. (*c*) Those projections averaged according to brain system.

**FIG. S11.**
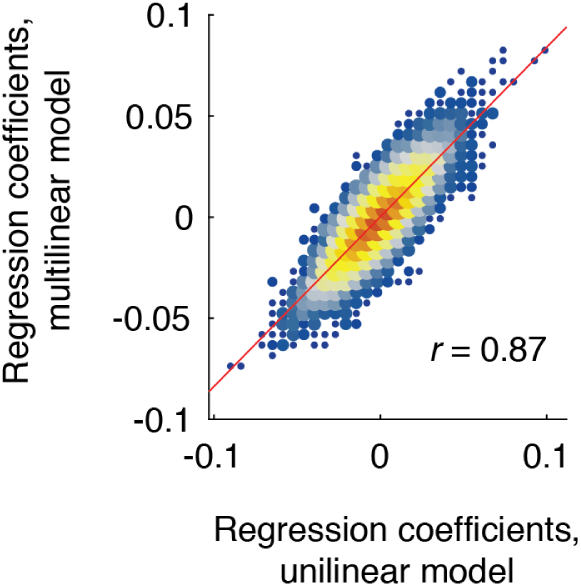
Comparison of regression coefficients estimated using uni-/multi-linear modeling. In the main text we reported *β* coefficients estimated from linear models of the form *ŷ* = *β*_0_ + *β*_1_*x*_1_ + *ϵ*. Here, we compare them against multi-linear models in which we allow many predictors to compete for the same variance. These models have the form *ŷ* = *β*_0_ + ∑_*i*_ *β*_*i*_*x*_*i*_ + *ϵ*. As means of comparison, we aggregated into a vector the coefficients for a given predictor (movie feature) for all connections. We repeated this procedure for both the uni-linear and multi-linear models and computed the correlation of those two vectors. In general, we find excellent correspondence. When all features are combined, we obtain a correlation of *r* = 0.87. Individually, there was more variability across predictors, but still with excellent correspondence, with mean ± standard deviation correlations of *r* = 0.88 ± 0.08.

**FIG. S12.**
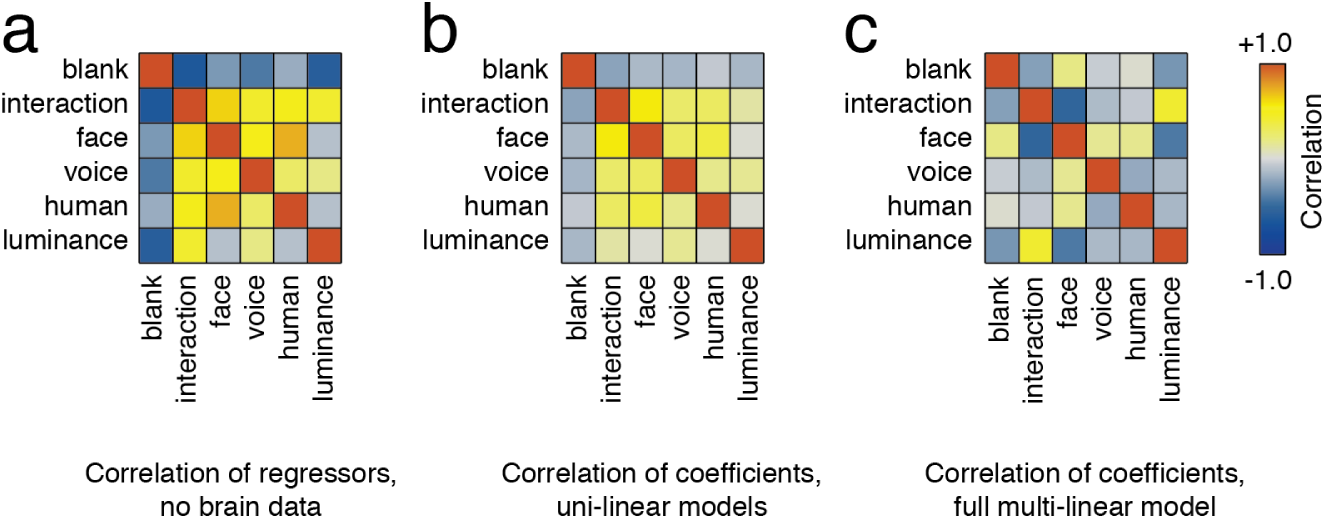
Impact of correlated regressors. In the main text we reported *β* coefficients estimated from uni-linear models of the form ŷ= *β*_0_ + *β*_1_*x*_1_ + *ϵ*. In Fig. S11, we compared them against multi-linear models in which we allow many predictors to compete for the same variance. Here, we show the correlation structure of *β* coefficients for both models alongside the correlation structure of the regressors, corresponding to a blank screen, an interaction between humans, presence of a face, voice, presence of a human, or baseline fluctuations in luminance. In general, we find that the regression structure of uni-linear models closely matches that of the regressors, themselves (*r* = 0.95). The multi-linear model, on the other hand, allows regressors to compete for the same variance, and as a result exhibits correlation structure dissimilar from the regressors on their own (*r* = − 0.06). We note, however, that the *β* coefficients from both the uni-linear and multi-linear models are, nonetheless, highly correlated (See Fig. S11).

**TABLE S1.** Movies included in each movie scan.

| Scan | Title | Genre | Runtime |
| --- | --- | --- | --- |
| 1 | Man Up and Go | documentary/emotional | 4m20s |
| 1 | The First 70 | documentary | 3m |
| 1 | Fixation | documentary/adventure | 1m42s |
| 1 | The Living | drama | 2m |
| 1 | SAMSARA | documentary/“unparalleled sensory experience” | 1m35s |
| 1 | Blood Brother | documentary | 2m20s |
| 2 | Birdmen | documentary/adventure | 3m59s |
| 2 | Groomed | drama | 1m30s |
| 2 | Cold | outdoor/sports | 2m |
| 2 | Sleepwalkers | drama | 2m |
| 2 | A Kind of Show | comedy | 1m |
| 3 | Geofish | documentary/adventure | 4m40s |
| 3 | The Debut | outdoor/sports | 3m23s |
| 3 | Dreams of a Life | documentary/mystery | 2m10s |
| 3 | The Front Man | documentary | 2m30s |
| 3 | This Is Vanity | drama | 1m |
| 4 | Planetary | documentary | 4m30s |
| 4 | Sign Painters | documentary | 2m50s |
| 4 | Florida Man | documentary/drama | 2m |
| 4 | The Sleeping Bear | drama | 3m40 |

1 For this analysis we truncated the fMRI BOLD time series for each session so that FC was always estimated using the same number of observations.

2 We note that the precise number and duration of these periods will depend upon the choice of statistical threshold.

